# Decellularization of Rat Gracilis Muscle Flap as a Potential Scaffold For Skeletal Muscle Composite Allotransplantation

**DOI:** 10.1101/2024.03.28.587269

**Authors:** Chenhui Dong, Aida K. Sarcon, Chunfeng Zhao

**Affiliations:** Biomechanics & Tendon and Soft Tissue Biology Laboratories, Division of Orthopedic Research, Mayo Clinic, Rochester, MN 55905, USA; Department of Sports Medicine, The 940^TH^ Hospital of Joint Logistics Support Force of PLA, Lanzhou Gansu 730050, China; Department of Surgery, Mayo Clinic, 200 First Street SW, Rochester, MN, USA

**Keywords:** decellularization, muscle, scaffold, biomaterial, allotransplantation

## Abstract

There are limited biomaterials for skeletal muscle regeneration. This study aimed to apply a decellularization protocol in a muscle flap model and investigate its patency. Twenty-six gracilis-muscle (GM) flaps were harvested from 13 rats. GMs were divided into groups of either 1) normal (control), 2) perfusion with 1% sodium dodecyl sulfate or SDS for 48h, followed by Triton X-100 or TX100, or lastly, 3) perfusion with SDS for 72h, followed by TX100. The morphology, microcirculatory network patency, and residual DNA content (DNAC) were evaluated. Decellularized muscle (DM) for 72h was more translucent than DM-48h. Despite longer decellularization, the DM-72h microcirculatory network maintained its integrity, except when the dye infiltrated from the muscle edges. Compared to normal, all DM had significantly lower DNAC (normal of 1.44 μg/mg *vs*. DM-48h of 0.37 μg/mg *vs*. DM-72h of 0.089 μg/mg; P < 0.001). The DNAC of the DM-72h group was significantly lower than DM-48h (P< 0.001). We report successful GM flap decellularization. Longer decellularization led to lower DNAC, which did not compromise circulation. Our protocol may be applicable as a free-flap scaffold model for transplantation in the future.

**Statement of clinical significance:** The impact of our work involves a reproducible skeletal muscle decellularization protocol to later apply in translational research.

## INTRODUCTION

Bioengineering is an emerging field in medicine. However, product feasibility is paramount and should be considered in product design. In the presence of devastating tissue loss, performing large donor tissue harvests may become impractical. This is important with a skeletal muscle injury, usually after trauma (i.e., combat injuries) or even after surgery (i.e., debridement of necrotizing soft tissue infections and/or extensive sarcoma resections). ^[1, 2]^ There has been growing interest in tissue engineering, and various options exist. One involves the use of stem cells, and this option has the theoretical advantage of cellular regeneration of the tissue of interest based on the microenvironment. However, studies have shown inconsistencies due to poor cellular retention and survival following inosculation.^[3]^

Another option is direct tissue transplantation from the patient (autotransplant) or other biomaterials (allotransplant). Historically, tissue auto or allotransplantation originated as early as the 1960s.^[4], [5]^ The surgical approach may change based on the tissue needed. Autologous tissue can be obtained locally, rearranged, or harvested from a distant donor site and then transplanted to the recipient site. A free muscle flap can fill a significant skeletal muscle defect. The first successful report of free skeletal muscle transplantation occurred in 1970 by Tamai et al.^[6]^ In a canine preclinical model, the authors transplanted a rectus femoris muscle onto a defect, and later the animal was able to achieve functional recovery with near complete histological evidence of muscle regeneration at five months postoperatively.^[6], [7]^ However, standard postoperative complications can still threaten any flap’s survival, which needs to be weighed against the initial donor site morbidity, and further flap reconstruction options after such a devastating complication. This may not always be a practical solution, particularly for large defects. In this regard, allografts are an attractive alternative as they negate the need for initial tissue harvest. However, autografts have excellent success rates in practice since they are entirely native to the patient.^[8], [9]^ Thus, another feasible option needs to overcome such barriers.

Another option is to bioengineer a scaffold that is tissue-specific and not immunogenic. Ideally, this would be an off-the-shelf product without further preparation before transplantation. There are several methods to engineer such scaffolds. One involves using naturally occurring materials such as tissue-specific extracellular matrix or ECM. ^[10]^These scaffolds would have the appropriate framework to fill the void, which can later be recellularized with the patient’s cells.^[11]^ Tissue decellularization is one method to create an ECM scaffold, as it allows the removal of cells and antigens.^[12], [13]^ Bioscaffolds have several applications, from solid organs to tissue valves and free-flaps. There are currently several bioscaffolds available in the market. These scaffolds include the heart,^[14]^ vessels,^[15]^ cartilage,^[16]^ and skin.^[17]^ Unlike these scaffolds, there is no Food and Drug Administration (FDA) approved skeletal muscle tissue bioscaffold.

Decellularization can be performed in several ways, either physically, chemically, or biologically with solutions to remove all cells and nuclear material.^[12]^ There is no standardized protocol,^[18]^ and an optimal protocol that maintains both the ECM integrity and circulatory integrity is needed for a successful scaffold free-flap transfer. For instance, hypertonic solutions may be used for decellularization and they are intended to cause osmotic shock and thus disrupt cellular bonds. ^[19]^ However, this may not entirely remove all of the cellular material.^[19]^ Additionally, the use of nucleases may result in an immunogenic response limiting its clinical utility in practice.^[19]^ Thus, the type and duration of each reagent used require careful review to preserve the scaffold’s ECM and eliminate cells that could be immunogenic. Our prior work used a detergent/perfusion and/or agitation protocol to decellularize a superficial inferior epigastric fasciocutaneous free flap.^[20]^ However, we could not achieve a “closed” or arterio-venous (AV) loop circulation. This study aimed to decellularize a free gracilis muscle (GM) flap using a detergent and perfusion protocol and later assessed the flap’s circulatory networks’ preservation.

## METHODS AND MATERIALS

### Gracilis Flap Harvest

Thirteen male Sprague-Dawley rats (12 weeks old, weighing 360 to 400 g) were used for this study. The GM was collected postmortem from humanely euthanized animals under an approved IACUC protocol at our institution. The GM was chosen since it is often used in clinical practice as a workhorse flap due to its versatility and low donor site morbidity.^[21]^ It is also a reliable flap rodent model.^[22]^ Male rats were selected due to their larger muscle mass which would facilitate easier harvest.

The rat was placed in the supine position with bilateral hindlimbs prepped. A curved incision was made just above the inguinal ligament on the limb, and dissection was taken until the GM was identified (Figure 1). The GM was dissected free with a femoral artery and vein. The GM flap was roughly 4cm x 3cm in size. The contralateral GM flap was harvested in the same fashion, yielding a total of 26 flaps.

**Figure 1A-D:**
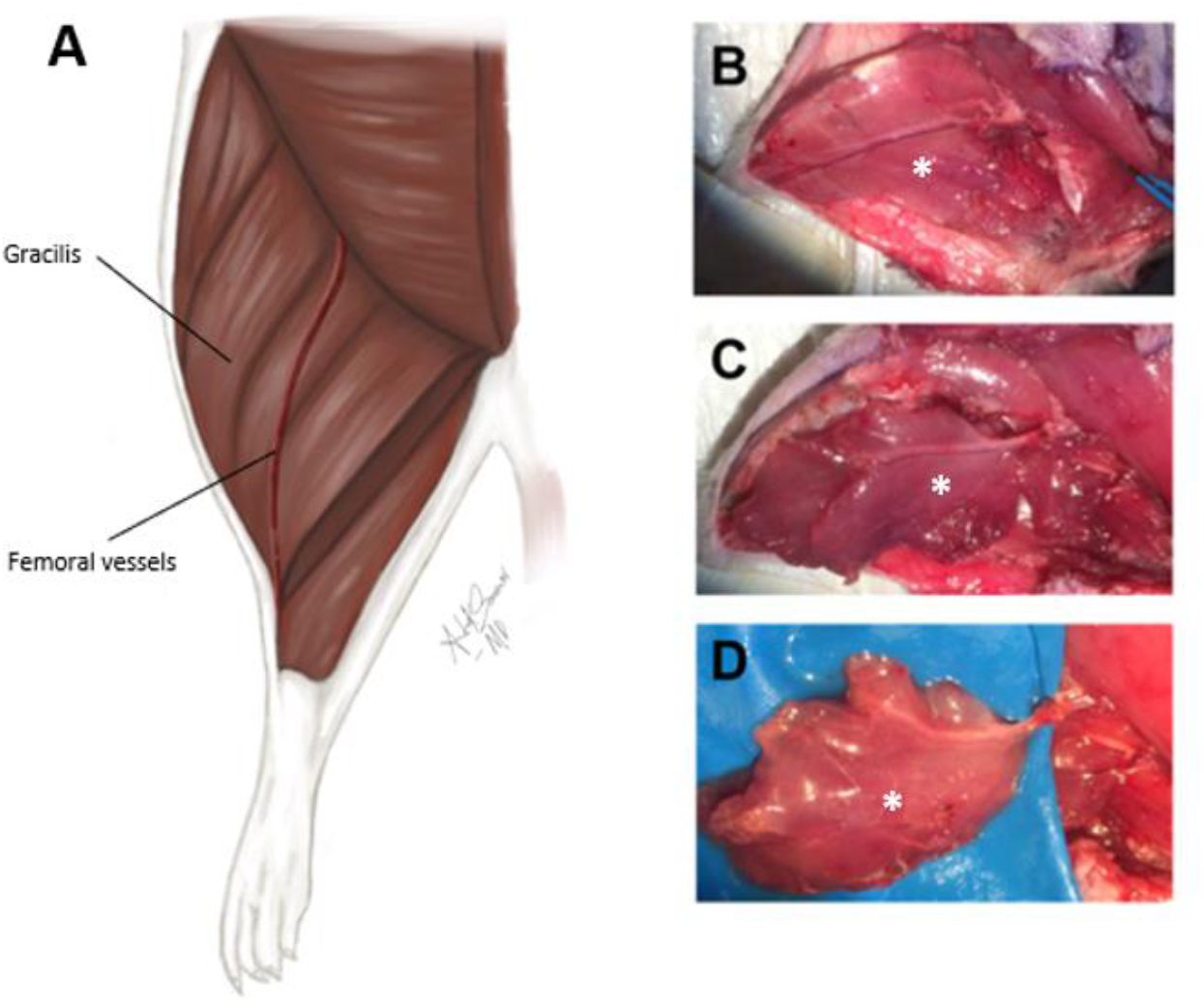
The gracilis muscle flap is harvested for decellularization. **A)** Illustration of the gracilis muscle with respect to the other hindlimb muscles and femoral vessels. **B-D)** Dissection and isolation of the gracilis muscle (asterisk) as seen with its vascular pedicle.

### Decellularization Protocol

Our decellularization protocol was adapted from our prior study, except for agitation alone.^[20]^ The GM flaps were each randomized into three groups, as seen in Figure 2. Briefly, the femoral artery was entered via a metal cannula and a universal pulmonary artery canulation system (Harvard Apparatus Holliston, MA) and this was secured using a nylon suture. All GMs were perfused per our study protocol (Figure 2) at a rate of 0.5 mL/min via a Syringe Pump (Fusion 100, Chemyx, Stafford, TX, USA) or Volumetric infusion pump (Baxter, Deerfield, IL, USA). Our prior experience predetermined this perfusion rate.^[20]^A total of six GMs were allocated into the normal or control group, which were only subjected to perfusion with heparinized phosphate-buffered saline (PBS). The experimental groups (decellularized muscle or DM) were also perfused with heparinized PBS. However, depending on the duration of the SDS, they were simultaneously perfused and soaked for either 48h (n=10) or 72h (n=10) with 1% SDS (Sigma-Aldrich®, USA). All GMs were then agitated with distilled water for 15 minutes, followed by perfusion and soaking with 1% Triton X-100 or TX100 for 30 min. Lastly, all were perfused and washed with PBS for 60 min.

**Figure 2:**
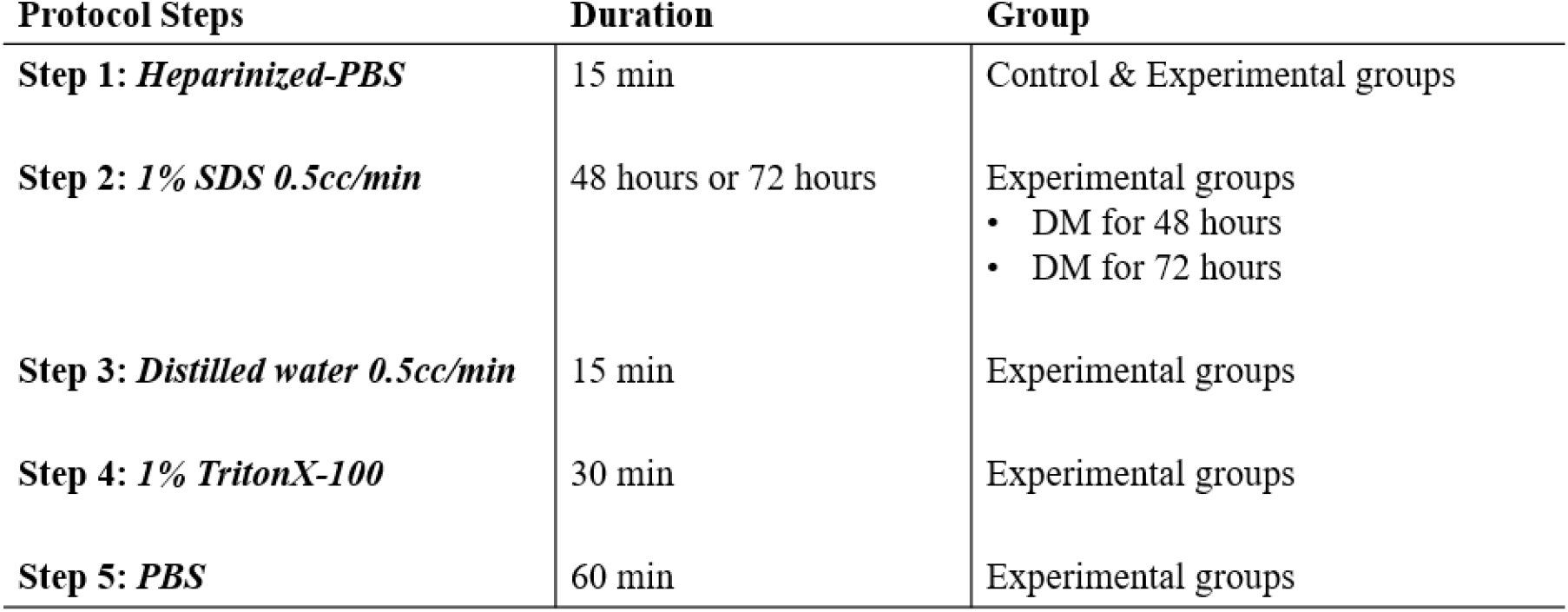
Flow chart of the decellularization protocol. Adapted from Qu *et al*. ^[20]^ Abbreviations: sodium dodecyl sulfate (SDS); Phosphate-buffered Saline (PBS)

### Arterial-Venous Circulation Patency and Microcirculatory Network Integrity

#### Arterial canulation

A total of n=2 GM flaps were chosen from each group to evaluate a patent femoral AV circulation and microcirculatory network. Due to its larger diameter, the femoral artery and vein were used as the desired flap pedicle. The femoral artery was entered with a metal cannula and attached to an intravascular polyethylene tube (Pulmonary artery ball joint system, Harvard Apparatus Holliston, MA) and later secured with a 6-0 polypropylene suture.

#### Injection of Trypan Blue Dye

Trypan Blue Dye (TBD), 0.4% solution, at a 0.5 mL/min perfusion rate, was used in this study like our prior published work.^19^ The perfusion video was recorded with the K-5II camera. The normal GM flaps were evaluated after heparinization since they were not subjected to decellularization. Following decellularization, the decellularized muscles in each group were immediately perfused with 0.4% TBD solution, and gross morphology was assessed for the integrity of the microcirculatory network.

### Gross Morphology

After undergoing the decellularization protocol, the GMs appearance was captured (K-5 IIs, Ricoh Company, Ltd. Japan). The muscles were visually inspected for their differences in their morphological appearance.

### Histology

A total of 2 GM flaps were chosen from each group for histological analysis. The muscles were fixed in 4% paraformaldehyde for 2 hours (Ratio of 20:1 fixative to tissue), and later paraffin-embedded. A 5-µm section was cut and stained with Hematoxylin-Eosin according to standard procedure. The slides were reviewed under microscopy (Olympus, BH-2, Tokyo, Japan) to evaluate the presence of skeletal muscle ECM and residual cellular material.

### DNA Quantification

A total of 8 GM flaps were used for DNA Quantification. The flaps were lyophilized for 72h with a freeze-drying machine (Millrock technology, Kingston, NY). A total of 25 mg of muscle weight was used to assess DNA content. Since the lymphatics/lymph nodes were found predominantly in the proximally inguinal region along the femoral artery, it was felt that the DNA content would be variable due to the presence of lymph nodes and, consequently, cells. Therefore, a region distally along the vascular pedicle was chosen to evaluate the residual DNA content for consistency. DNA was extracted per manufacture protocol using Qiagen DNeasy Blood and Tissue Kit (Valencia, CA, USA). The DNA concentration was assessed per manufacture protocol using Quant-iT PicoGreen dsDNA assay kit (Invitrogen, Carlsbad, CA). Fluorescence was measured using a microplate reader (FLUOstar Omega, BMG LABTECH, and Ortenberg, Germany). The DNA concentration (or content) was obtained from a standard curve.

### Statistical Analysis

The quantitative residual DNA data was represented as averages with standard deviation. Data were analyzed using the *t*-test with a significance set at an alpha of 0.05.

## RESULTS

### Longer Decellularized Muscle Is More Translucent

The gross evaluation of the GM is seen in Figure 3A-D. The GM is seen after perfusion with heparinized PBS (Figure 3A). The artery and vein were apparent in the GM flap; however, the tissue became more translucent with increasing duration of SDS (Figure 3B to D). The DM 48h had patchy translucent areas compared to the nearly complete translucent DM 72H (Figure 3C). The most translucent DM followed agitation with distilled water for 30 minutes at a constant rate, followed by 1% Triton X-100 perfusion and later 60 minutes PBS perfusion with soaking (Figure 3D).

**Figure:3A-D.**
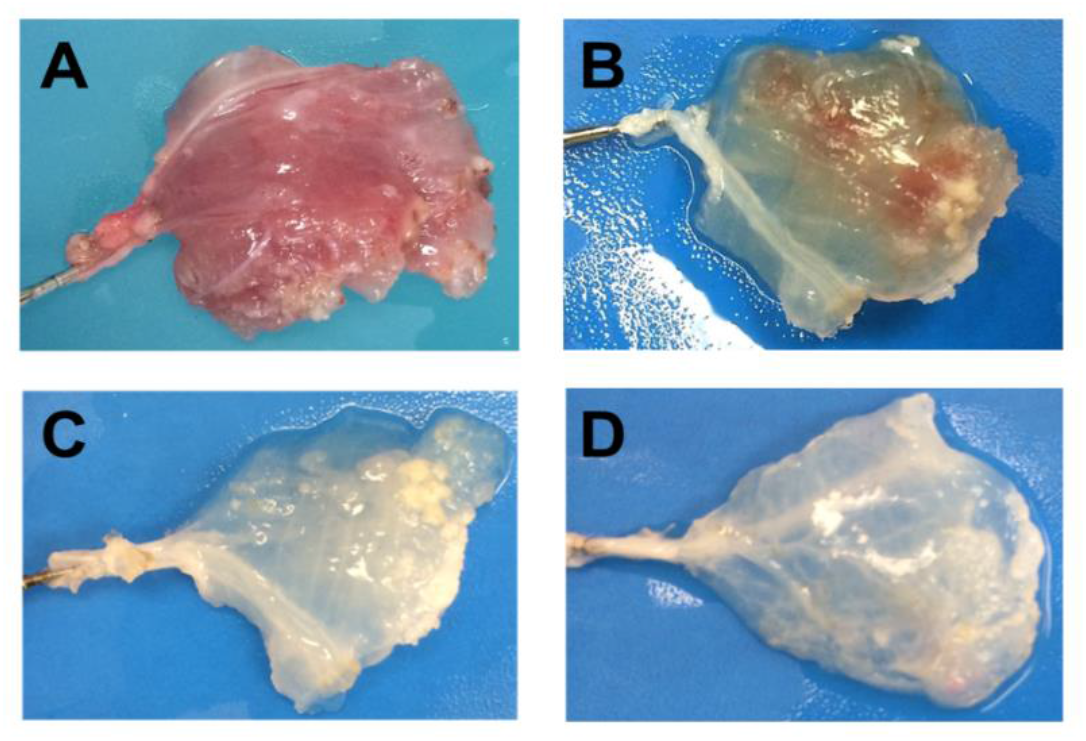
Gross morphological evaluation of the gracilis muscle (GM) flap per study protocol. **A)** demonstrating harvested control group GM flap compared to decellularization of muscle (DM) as seen on B-D; **B)** GM perfused with 1% SDS for 48h resulting in patchy translucency compared to **C)** which is near complete translucency with DM 72h. **D)** Complete translucent DM after agitation with distilled water for 15 min and later perfusion and soaking with Triton X-100, followed by PBS.

### Longer Decellularized Muscle Has Less Retained Cells

The histological findings are shown (Figures 4 and 5). The normal muscle group showed intact myofibers surrounded by perimysium (Figure 4A&D). In contrast, DM 72h had decellularized muscle with only the ECM remaining (Figure 4C &F). A similar trend was seen with other cell structures (Figure 5). The control group had preserved epithelial cells, smooth muscle cells along the vessel walls, fibroblasts, red cells, lymphocytes, adipocytes, and endothelial cells (Figure 5A-D). Histologically, the number of cellular materials appeared to reduce with decellularization. The DM-48h group showed some residual cells along the artery, vein, and lymphatics (Figure 5E-H). In contrast, the DM 72h had none remaining (Figure 5I-L).

**Figure 4A-F:**
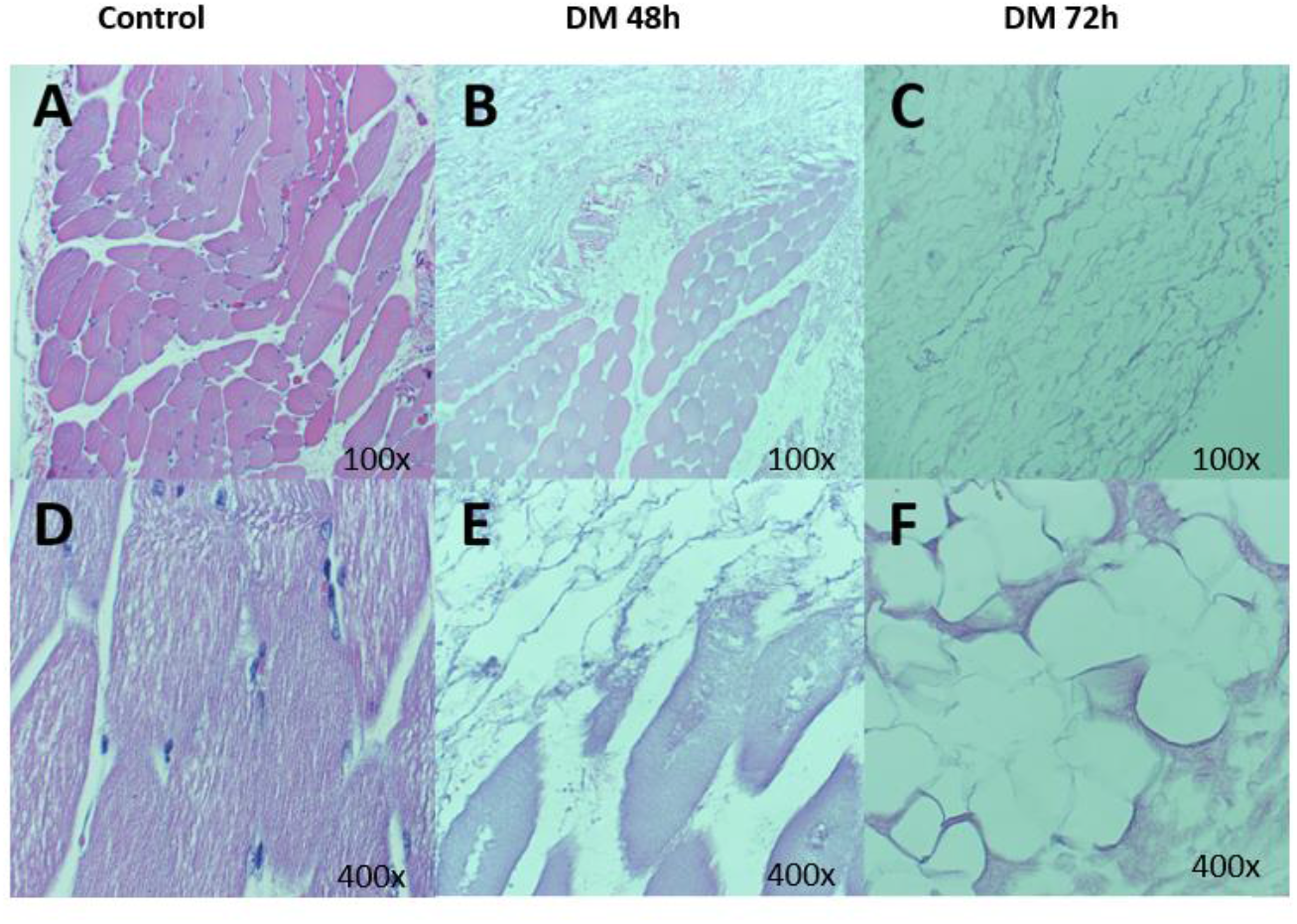
Histological H&E evaluation of muscle architecture in the study groups. The image demonstrates the remaining myofibers and myocytes. **A&D** control muscles with intact myofibers and remnant myocytes; **B & E)** 48 h decellularization of gracilis muscle (GM) resulted in reduced myofibers and myocytes, compared to **C &F)** 72h decellularization, which shows no remaining cells.

**Figure 5A-L:**
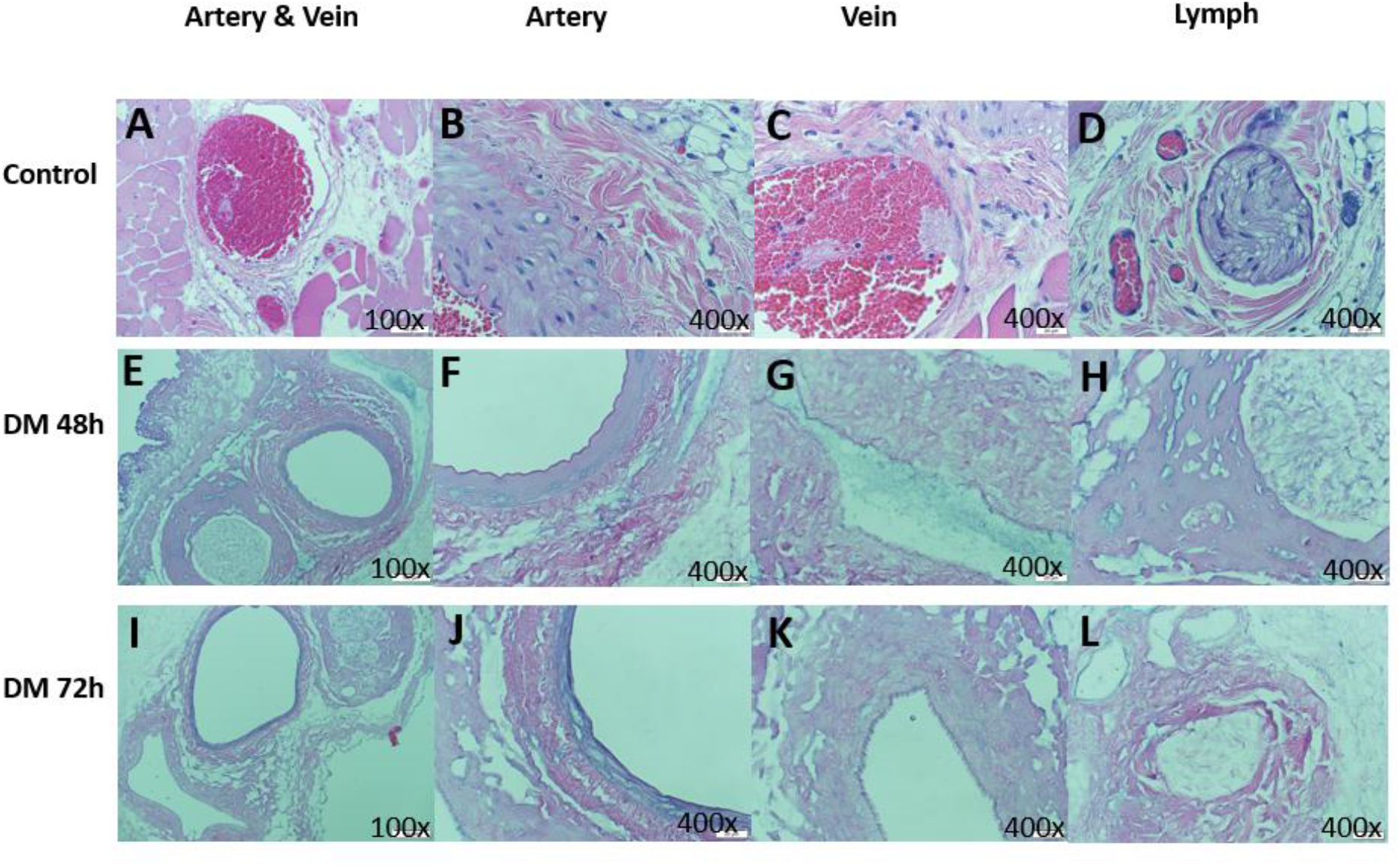
Histological H&E evaluation of arterio-venous and lymphatic structures. **A-D** control tissue shows intact cellular framework artery, vein, and lymph; **E-H)** following 48 h decellularization, there are residual cells along the artery, vein, and lymphatics **C&F)** 72h decellularization resulted in no cellular material.

### Decellularized Muscle Has a Patent circulation

The integrity of the microcirculatory network of the GM flap is shown in Figure 6. Compared to the control, the overall size of the DM 72hr was smaller and thinner by gross visual observation (Figure 6A). When the DM-72h group was perfused with the TBD dye, the dye immediately distributed into the flap’s vascular network; most of it returned from the accompanying vein, some dye stayed in the flap, and only sometimes did the dye leak from the flap edges (supplemental video). The microcirculatory network appeared intact after our 72h decellularization protocol (Figure 6B).

**Figure 6A-B:**
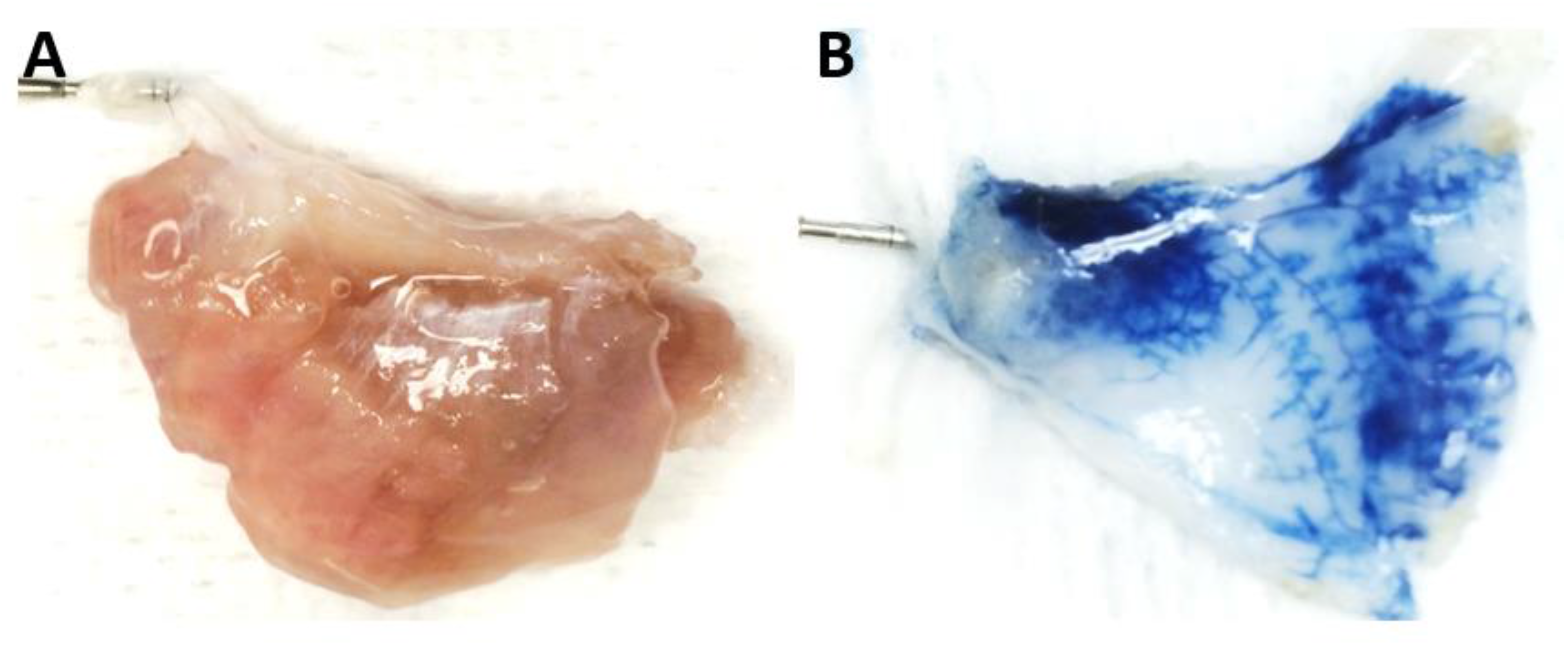
Gross morphology of the gracilis muscle (GM) flap. **A)** normal or control GM, and **B)** 0.4% Trypan blue dye blue was immediately used to perfuse the vascular pedicle to confirm microcirculatory network integrity in decellularized GM 72h.

### Longer Decellularized Muscle Has Lowest Remaining DNA

The DNA content of the groups is shown (Figure 7). Compared to the control group (1.44µg/mg), there was significantly less DNA in both DM groups (P < 0.001). There DM 72h had lower DNA content compared to DM 48h (0.089µg/mg *vs* 0.38 µg/mg; P < 0.001).

**Figure 7:**
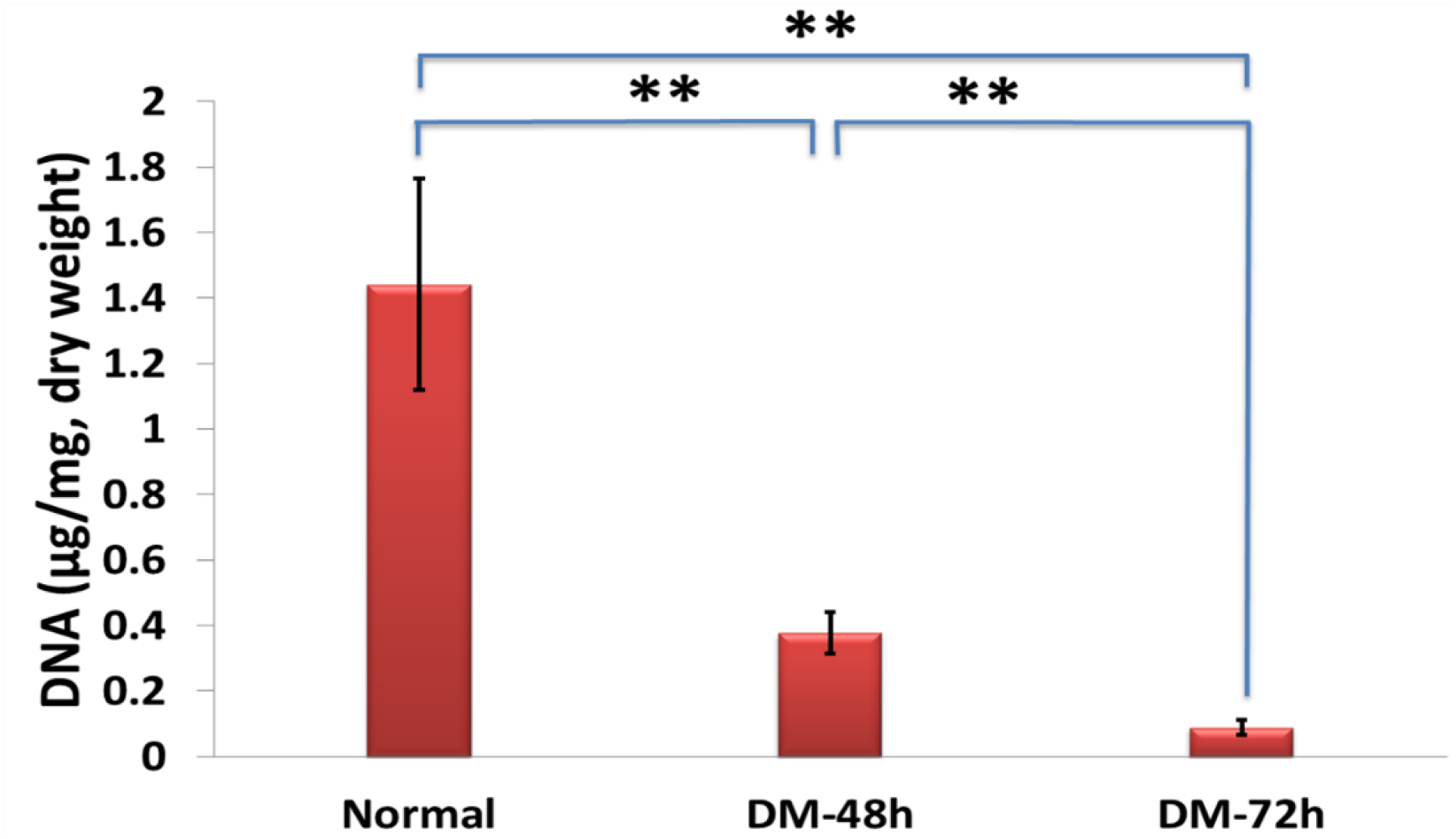
All decellularized muscles (DM) showed lower DNA content compared to the control. (1.44µg/mg, P<0.001). The amount of DNA content significantly reduced with increasing decellularization (DM 72h of 0.089µg/mg*vs* DM 48h 0.38 µg/mg, p <0.001)

## DISCUSSION

Several bioscaffolds in the market have been used for tissue transplantation. One of its theoretical benefits is the low immunogenicity (due to the fewer antigens), which negates the need for immunosuppressive therapy postoperatively. However, there is still no FDA-approved skeletal muscle scaffold on the market. This can be partially attributed to its complex cellular regeneration process.^[23-25]^ A free muscle flap is commonly used in clinical practice and has many applications for soft tissue coverage.^[26], [27]^ However, it carries the risk of donor site morbidity and thus needs to be thoughtfully considered during surgical planning. A skeletal muscle bioscaffold could overcome this barrier as it would avoid the need for donor site harvest. In our study, we decellularized a GM flap, resulting in translucent morphology with less cellular content and residual DNA. If dye leakage occurred, this was at the transected muscle edges. To the best of our knowledge, in the current study, we, for the first time, employed this perfusion technique to decellularize the muscle flap.

The significant finding of our study was our ability to maintain patent circulation, which was a substantial barrier in our prior lab’s work. In our previous study, we decellularized a superficial epigastric fasciocutaneous flap using either detergent perfusion and/or perfusion decellularization method, and we found that the combination technique was superior to agitation alone due to less residual DNA.^[20]^ However, none of our flaps could maintain a “close loop” circulation. One explanation for this finding could be due to the nature of the flap harvested in our prior study and the caliber of the vascular pedicle to allow for perfusion and venous return. After decellularization, we found that most of the dye leaked from the vein and some from the actual flap. In another flap model, the authors demonstrated an intact AV circulation after 6 h of perfusion of a decellularized flap embedded in agarose with a peristaltic pump at a 0.2 ml/min rate.^[28]^ We chose a rate of 0.5 ml/min rate to perfuse the GM via a peristaltic perfusion pump which provided the ability to preserve the microcirculatory network and ECM.

There are several decellularization protocols with differences in duration and solution. Sabbagh *et al*. also performed decellularization of the GM flap.^[29]^ The authors evaluated two different DM groups; one group underwent detergent/perfusion with Krebs Henseleit buffer (KHB), 1% SDS 1h and 1% TX100 1h, and the other with 1% SDS for 72 hr, 1% TX100 1h. They found patent circulation in both DM groups. Concerning histology, they discovered that the 3D architecture was less distorted with the KHB group.^[29]^ The authors found a statistically significant reduction in DNA content irrespective of the assigned DM group. Similarly, we found a significant decrease in DNA content regardless of time, although with the most significant decrease with DM 72h. Additionally, this translated to a more translucent morphology since DM 72 h was clearer than the other groups. This suggests that using the flap’s pedicle provided reliable access and a method to perform our decellularization protocol.^[19]^

The current study has several limitations. First, the perfusion pressure was not monitored or controlled. However, the pressure could be inferred from prior studies that used similar perfusion rates within this range. Studies have found that a rate of 0.2 ml/min in a similar flap model corresponds to a perfusion pressure of around 60 to 120 mmHg.^[28], [30]^ A recent review article showed that increased pressure may not have detrimental effects on a vessel if the perfusion is maintained throughout.^[19]^ However, future research on this topic may help optimize specific perfusion pressures, rates, and effects on DM. The integrity of the microcirculatory network was only observed through qualitative dye infusion without quantifying the vascular network. The DNA content was also obtained from a select region of the tissue (distal to the vascular pedicle); thus, the values are not representative of the total residual DNA in the flap. Lastly, we did not evaluate different concentrations of SDS in the decellularization process as it was felt to be outside of the aim of the present study. We also did not evaluate DM at shorter durations, and we may have reached a statistically significant reduction of DNA earlier. Future research on this matter would be helpful in further optimizing SDS detergent-mediated decellularization protocols.

## CONCLUSIONS

We report a reproducible protocol to decellularize a GM flap using 1% SDS for either 48 or 72 h duration, followed by 1% TX100. The flap maintained a viable circulatory network despite lower DNA content and complete translucent structure after 72 hr of decellularization. This is important when designing a free flap scaffold for transplant to a recipient site. A future study that evaluates the application of such a decellularized scaffold in an animal model incorporating recellularized stem cells is warranted and will shed further insight into this protocol.

## SUPPLEMENTARY MATERIALS

Video of perfusion test

## ACKNOWLEDGEMENTS

none

## DATA AVAILABILITY STATEMENT

The data presented in this study are available upon reasonable request from the corresponding author.

## REFERENCES

1. Dawson JI, Kanczler J, Tare R, et al. Concise review: bridging the gap: bone regeneration using skeletal stem cell-based strategies - where are we now? Stem Cells. 2014; 32(1):35–44.

2. Cheng, CS, Davis BD, Madden L, et al. Physiology and metabolism of tissue-engineered skeletal muscle. Exp Biol Med (Maywood). 2014; 239(9): 1203–14.

3. Ostrovidov S, Hosseini V, Ahadian S, et al. Skeletal muscle tissue engineering: methods to form skeletal myotubes and their applications. Tissue Eng Part B Rev. 2014;20(5):403–36.

4. Lambert PB, & Frank HA. Local recognition of histocompatibility differences in skin grafts. Science.1967;155(3758):99–101.

5. Irish JD. A 5,500 year old artificial human tooth from Egypt: a historical note. Int J Oral Maxillofac Implants.2004;19(5):645–7.

6. Tamai S, Komatsu S, Sano S, et al. Free muscle transplants in dogs, with microsurgical neurovascular anastomoses. Plast Reconstr Surg.1970: 46(3):219–25.

7. Harii K. Free Skeletal Muscle Transfer With Microneurovascular Anastomoses. In: Muscle Transplantation. (Freilinger, G. Holle J, Carlson BM. eds.) Springer, Vienna.1981.

8. Cooper J, & Beck C. History of Soft-Tissue Allografts in Orthopedics. Sports Medicine and Arthroscopy Review.1993;1(1): 2–16.

9. Simsiman AJ, Powell CR, Stratford RR, et al. Suburethral sling materials: best outcome with autologous tissue. Am J Obstet Gynecol.2005;193(6):2112–6.

10. Wolf MT, Dearth CL, Sonnenberh SB, et al. Naturally derived and synthetic scaffolds for skeletal muscle reconstruction. Adv Drug Deliv Rev. 2015;84:208–21.

11. Hillebrandt KH, Everwien H, Haep N, et al. Strategies based on organ decellularization and recellularization. Transpl Int.2019;32(6): 571–585.

12. Srokowski EM, Woodhouse, KA. 2.20 Decellularized Scaffolds. In Comprehensive Biomaterials II (Ducheyne P. eds.) Elsevier. 2017. 452–470.

13. Macchiarini P, Jungebluth P, Go T, et al. Clinical transplantation of a tissue-engineered airway. Lancet. 2008; 372(9655): 2023–30.

14. CardioCel BioScaffoled. 2022. Available from: https://www.lemaitre.com/products/cardiocel-bioscaffold. [Last accessed 12/20/2022]

15. VascuCel Bioscaffold. 2023. Available from: https://www.lemaitre.com/products/vascucel-bioscaffold. [Last accessed 12/20/22]

16. Hede K, Christensen BB, Olesen ML, et al. CARGEL Bioscaffold improves cartilage repair tissue after bone marrow stimulation in a minipig model. J Exp Orthop. 2020; 7(1): 26.

17. PriMatrix® Dermal Repair Scaffold. 2023. Available from: https://www.integralife.com/primatrix-dermal-repair-scaffold/product/wound-reconstruction-care-inpatient-acute-or-primatrix-dermal-repair-scaffold. [Last accessed 12/20/22]

18. Reyna WE, Pichika R, Lidvig D, et al. Efficiency of skeletal muscle decellularization methods and their effects on the extracellular matrix. J Biomech. 2020;110: 109961.

19. Crapo PM, Gilbert TW, & Badylak SF. An overview of tissue and whole organ decellularization processes. Biomaterials. 2011;32(12):3233–43.

20. Qu J, Van Hogezand RM, Zhao C, et al. Decellularization of a Fasciocutaneous Flap for Use as a Perfusable Scaffold. Ann Plast Surg. 2015; 75(1):112–6.

21. Lyons ME and Goldman. Gracilis Tissue Transfer. In StatPearls. StatPearls Publishing. Treasure Island FL. 2024

22. Dautel G, Braga da Silva JL, and Merle M. Pedicled or Free Flap Transfer of the Gracilis Muscle in Rats. Journal of Reconstructive Microsurgery. 1991; 7: 23–25

23. Yao KA, Roth AC, Stephenson LL, et al. Effect of ketorolac tromethamine (Toradol) on ischemia-reperfusion injury in skeletal muscle. J Reconstr Microsurg. 1998; 14(3): 211–4.

24. Gillani S, Cao J, Suzuki T, et al. The effect of ischemia reperfusion injury on skeletal muscle. Injury. 2012; 43(6):670–5.

25. Garg K, Ward C, Rathbone C, et al. Transplantation of devitalized muscle scaffolds is insufficient for appreciable de novo muscle fiber regeneration after volumetric muscle loss injury. Cell Tissue Res. 2014;358(3):857–73

26. Lin CH, Lin YT, Yeh JT, et al. Free functioning muscle transfer for lower extremity posttraumatic composite structure and functional defect. Plast Reconstr Surg. 2007; 119(7): 2118–2126.

27. Stevanovic M, & Sharpe F. Functional free muscle transfer for upper extremity reconstruction. Plast Reconstr Surg. 2014;134(2):257e–274e.

28. Henderson PW, Nagineni VV, Harper A, et al. Development of an acellular bioengineered matrix with a dominant vascular pedicle. J Surg Res. 2010;164(1):1–5.

29. Sabbagh MD, Roh SG, Lui J, et al. A Quick and Reliable Method to Decellularize a Gracilis Flap: A Crucial Step Toward Building a Muscle. Ann Plast Surg. 2019; 83(6): 709–715.

30. Chang EI, Bonillas RG, El-ftesi S, et al. Tissue engineering using autologous microcirculatory beds as vascularized bioscaffolds. FASEB J. 2009;23(3): 906–15.

